# Network connectivity differences in music listening among older adults following a music-based intervention

**DOI:** 10.1101/2024.06.13.598944

**Authors:** Sarah Faber, Alexander Belden, Randy McIntosh, Psyche Loui

## Abstract

Music-based interventions are a common feature in long-term care with clinical reports highlighting music’s ability to engage individuals with complex diagnoses. While these findings are promising, normative findings from healthy controls are needed to disambiguate treatment effects unique to pathology and those seen in healthy aging. The present study examines brain network dynamics during music listening in a sample of healthy older adults before and after a music-based intervention. We found intervention effects from hidden Markov model-estimated fMRI network data. Following the intervention, participants demonstrated greater occupancy (the amount of time a network was occupied) in a temporal-mesolimbic network. We conclude that network dynamics in healthy older adults are sensitive to music-based interventions. We discuss these findings’ implications for future studies with individuals with neurodegeneration.

## Introduction

Listening to music is an enjoyable activity for younger and older adults and is a common feature in long-term care settings. Studies of music-based interventions offer promising glimpses into the efficacy of music as an intervention for clinical populations (see Särkämö et al., 2014; Cuddy & Duffin, 2005, Satoh et al., 2015; Guétin et al., 2009; Svansdottir & Snaedal, 2006). Few studies exist on music-based interventions in healthy populations, rendering clinical work findings difficult to contextualize. Further complicating the research landscape is heterogeneity in healthy adult populations with no clear consensus on how to predict the onset and trajectory of age-related pathologies, such as dementia.

As the brain ages, its organizational structure changes, culminating in network reconfigurations. At rest and during cognitive tasks, within-network functional connectivity (FC) in canonical networks (such as the default mode, salience, and dorsal attentional networks) is reduced in older adults compared to young adults, with corresponding increases in FC between networks (Grady et al., 2016). These reconfigurations describe dedifferentiation: a process where brain activity associated with specific cognitive networks (such as the default mode network) becomes more integrated across the brain and less functionally segregated (Grady et al., 2012).

Dedifferentiation has been seen in the default mode network (DMN), fronto-parietal network (FPN), and salience ventral attention network at rest (Malagurski et al., 2020). The activity of these networks also changes with age with older adults showing lower attenuation of the DMN and correspondingly lower activation in the dorsal attention network (DAN) during cognitive tasks. These organizational shifts start in middle age and do not predict task performance, which emerges much later in adulthood (Rieck et al., 2017). Between the networks, FC patterns become less distinct between the DMN, DAN, FPN, and the auditory subsection of the somatomotor network (Rieck et al., 2021) with the FPN becoming more connected to the DMN and DAN with age (Rieck et al., 2017). This is not a blanket effect across the brain: the ventral attention network and the motor subsection of the somatomotor network show increased within-network connectivity later in life (Grady et al., 2016; Rieck et al., 2021).

In dynamic FC studies differences between rest and task again emerge with older adults showing reduced within-network FC in the DMN during cognitive tasks compared to younger adults, but not at rest (Grady et al., 2016). Older adults also show higher occupancy in higher-order visual networks compared to younger adults in visual tasks (Tibon et al., 2020), indicating modality-specific aging effects.

While dedifferentiation-related changes in network organization are reliably seen with age, they may not consistently predict behavioural performance (Rieck et al., 2017), and vary between sensory modalities (Grady et al., 2016; Rieck et al., 2021). Certain reconfigurations may also contribute to resilience. In inhibitory processing and executive function tasks, greater integration between the DAN and other networks has been shown to correlate positively with task performance (Rieck et al., 2021; Reinberg et al., 2015).

In the auditory domain, age is commensurate with a decline in auditory acuity (Freiherr et al., 2013), but while hearing loss is associated with poor cognition (Lin et al. 2013), whether hearing loss leads to cognitive decline, or whether cognitive decline leads to degradation of the auditory system, remains unclear (Sardone et al., 2019). Parsing complex signals becomes more difficult with age (Alain et al., 2006), which has been studied with language (see Schneider et al., 2010) and music (Andrews et al., 1998), and may be mitigated by musical training (Zendel & Alain, 2012; Alain et al., 2014).

Musical training and listening can change the brain structurally. Older adults with a history of musical training and music listening show greater volume in the parahippocampus, and inferior frontal gyrus (specifically the pars opercularis and pars orbitalis); regions related to memory and language processing (Chaddock-Heyman et al., 2021). In healthy adults, music is a common tool for mood regulation (see Thayer et al., 1994; Saarikallio & Erkkilä, 2007). For those with Alzheimer’s disease, music listening and musical memory have been shown to engage a network of regions, including the ventral pre-supplementary motor area and caudal anterior cingulate gyrus, that show little atrophy and metabolic disruption, despite equivalent amyloid deposits with the rest of the brain (Jacobsen et al., 2015). Global FC increases immediately following preferred music listening (King et al., 2018), and familiar music listening in Alzheimer’s disease and mild cognitive impairment (MCI) is associated with greater activity in medial pre-frontal regions, the anterior insula, precuneus, basal ganglia, hippocampus, amygdala, and cerebellum compared to listening to novel music (Thaut et al., 2020). In healthy older adults, greater FC between auditory and reward regions has been observed following a regular music listening intervention (Quinci et al., 2022).

Behaviourally, individuals with mild and moderate Alzheimer’s disease perform on par with cognitively healthy peers in emotion recognition in music tasks (Arroyo-Annló et al., 2019) and identification of familiar melodies (Hsieh et al., 2011). A recent review of music-based interventions in Alzheimer’s care reported better outcomes for individually-selected music versus other-selected music in cognitive domains and for behavioural downregulation, while active interventions, such as singing, playing, etc., had better outcomes for behavioural upregulation (Leggieri et al., 2019).

Establishing how the brain network landscape changes with music-based interventions in healthy older adults will provide vital information for music and health research by establishing an age-appropriate normative baseline for future clinical work. In the present study, we use data from Quinci et al. (2022) to quantify brain network differences in a healthy older adult population before and after an 8-week music listening-based intervention. We aim to provide normative patterns of brain network activity for future comparative work with individuals with cognitive decline.

## Methods

### Participants

Older adults (age range 54-89, *N* = 27, 13 males, mean age = 67.34, *SD* = 8.27) were recruited for a music listening study through Northeastern University. They were right-handed and cognitively healthy and had normal hearing established via audiogram. Participants were excluded from the study if they had a change in medication within six weeks of screening, a history of cognitive impairments, a history of chemotherapy 10 years before screening, and any medical conditions requiring medical treatment three months prior to screening. The study complied with the Declaration of Helsinki and received ethics approval by the Northeastern University Institutional Review Board.

Participants were screened by researchers before data collection both to confirm their eligibility for the study and to collect a selection of familiar, well-liked songs to be presented during scanning. A pre-intervention scanning session included fMRI data collection, a blood draw, and a battery of neuropsychological tests. Participants then met with a board-certified music therapist (MT-BC) to develop personalized music playlists to be used during the intervention. Following completion of the intervention, participants returned for a post-intervention scanning session where they repeated the measures in the pre-intervention session. The present report details fMRI findings exclusively. Other measures will be detailed in future reports.

### Data Acquisition

Scanning occurred at Northeastern University. A Siemens Magnetom 3T scanner (64-channel head coil) was used to collect functional scans at a TR of 475 ms over 1440 volumes. Participants completed a resting state block followed by blocks of music listening. There were 24 musical excerpts, and each excerpt was either participant-selected and well-liked and familiar (6/24); or experimenter-selected. The experimenter-selected music fell into two sub-categories: songs that were from the popular canon during the participant’s youth (10/24), or songs that were selected to optimize novelty (8/24). Each excerpt was presented for 20 seconds in a random order and participants rated liking and familiarity on a 4-point Likert scale following each excerpt. Total scan time was 11.4 minutes.

### Data pre-processing

Functional MRI data were preprocessed using customized scripts from the TVB-UKBB pipeline (Frazier-Logue et al., 2022). MNI T1 templates were used for T1 image registration, and functional data preprocessing employed a sub-pipeline using tools from the FMRIB software library (Woolrich et al., 2009) to complete gradient echo field map distortion correction and motion correction (using MCFLIRT). We classified artifacts using independent component analysis (ICA) and, using MELODIC and FIX, assembled a training set for automatic artifact detection of the sample using the data from eight participants in the current study. Output was visually inspected by researchers for accuracy in artifact classification. Preprocessed datasets were downsampled to 220 regions using the Schaefer cortical and Tian sub-cortical atlases (Schaefer et al., 2018; Tian et al., 2020), and z-score normalized region timeseries data were exported to MatLab (MathWorks, 2019) for further analyses.

Twenty-four older adults passed MRI screening and completed pre-intervention scans. At the time of writing, eighteen participants had completed the intervention and post-intervention scans. Three participants were excluded from the final analysis for excessive head motion and/or problems with the behavioural data recording apparatus, resulting in a final sample of fifteen older adults (8 men, age range: 54-89 years, mean age = 62.67, SD = 15.35).

### Intervention and protocol

The music-based intervention (MBI) was a self-administered active listening music intervention based on work detailed by Hanser and Thompson (1994). Participants met with a board-certified music therapist (MT-BC) prior to the start of the intervention to collaboratively develop two personalized playlists: one of relaxing songs, and one of energizing songs. Each playlist was stored on a YouTube Premium account provided by the researchers and part of the MT-BC’s initial meeting with participants was to provide instruction in accessing and modifying the playlists. The MBI consisted of one hour of active listening to selections from one or both playlists each day for eight weeks. Participants were instructed to attend to any music-induced images, feelings, or memories; and to note if and how the music affected their mood during listening. Following listening, participants were asked to record their observations in a journal. Once weekly, participants would meet via Zoom with the MT-BC to review these observations, discuss the intervention, and adjust the playlists, if necessary.

Participants completed two fMRI scanning sessions, one prior to beginning the intervention, and the second following completion of the intervention. Task fMRI were the same stimuli described in the data acquisition section (six self-selected excerpts, and 18 experimenter-selected excerpts; 10 excerpts that were popular and/or recognizable, eight purpose-composed for research purposes), and were repeated during the post-intervention fMRI scanning session.

### State Estimation and Analysis

Brain state estimation was completed using the HMM-MAR toolbox (Vidaurre et al., 2017) and has previously been reported in Faber et al., 2023. A review of the estimation is presented here, with a re-print of the regions of interest (ROIs) comprising each estimated state.

We extracted two key features from the HMM model output for further analysis: fractional occupancy and transitional probability matrices. Fractional occupancy describes the number of time windows each state is occupied per target time window, and the transitional probability matrix describes the pairwise likelihood of transitioning from each state to each other state.

We used partial least squares (PLS) analysis to differentiate the combined pattern of HMM features that distinguished treatment effects. PLS is a multivariate technique that identifies latent patterns in matrices of manifest variables. These patterns are returned as latent variables (LVs), and bootstrap estimation and permutation testing are used to establish reliability and statistical significance for each LV (McIntosh & Lobaugh, 2004). We have chosen PLS over other similar methods (most notably, canonical correlation analysis) as it performs reliably with the presence of high within-block correlations (McIntosh, 2021).

## Results

### Fractional occupancy and transitional probabilities

We calculated fractional occupancy and transitional probability matrices for each participant during music listening for each category of musical stimulus (self-selected, experimenter-selected popular, and experimenter-selected novel) pre- and post-MBI. There was an intervention effect showing higher occupancy in the medial frontoparietal state (state1) and the temporal-mesolimbic state (state 3) for all music categories in the post-intervention condition (Figure 2).

**Figure 1:**
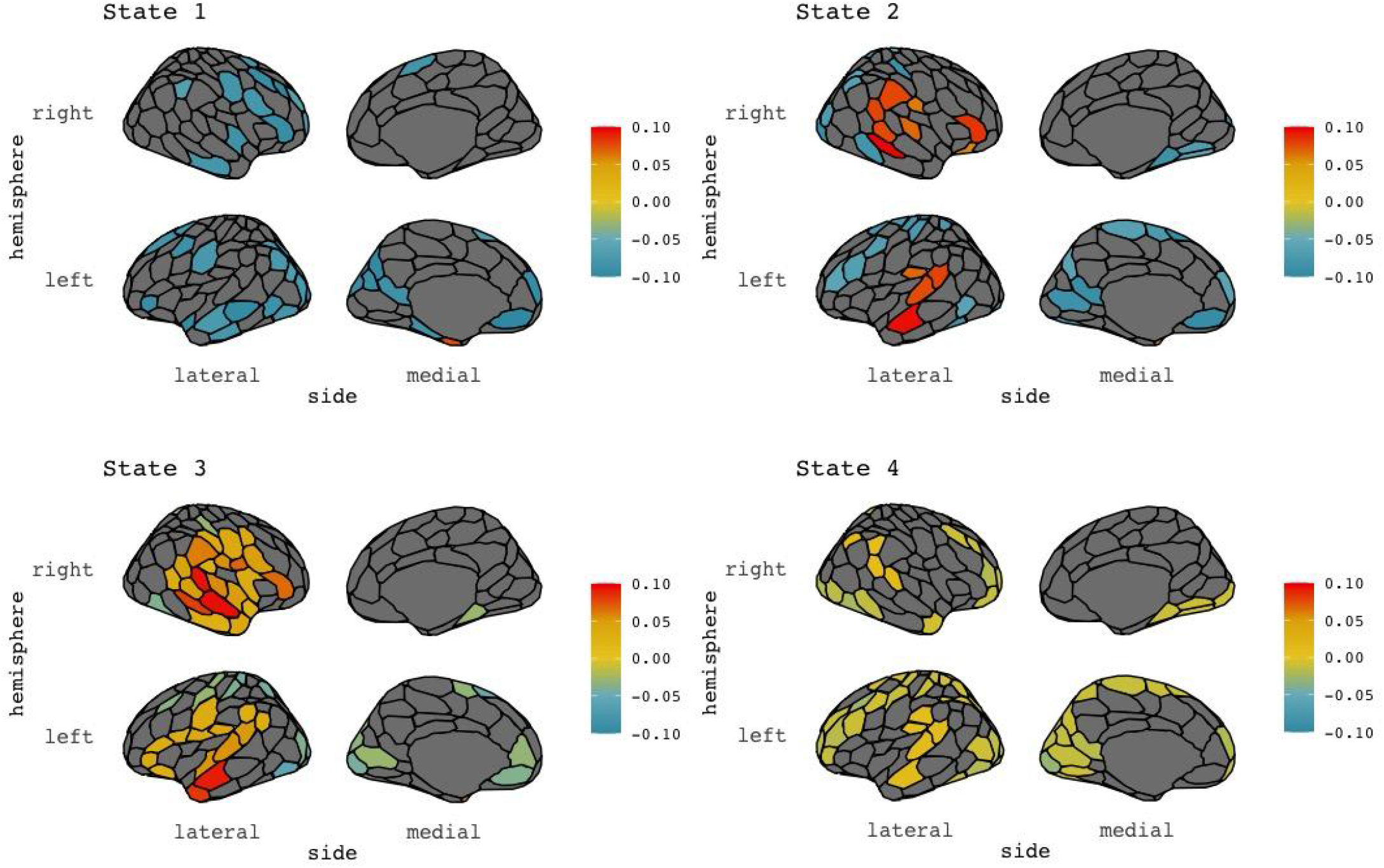
Mean activity plots returned from HMM analysis. Colours represent relative activity of the states and all have been normalized within-state. See Table 1 for subcortical regions not displayed here.

**Figure 2:**
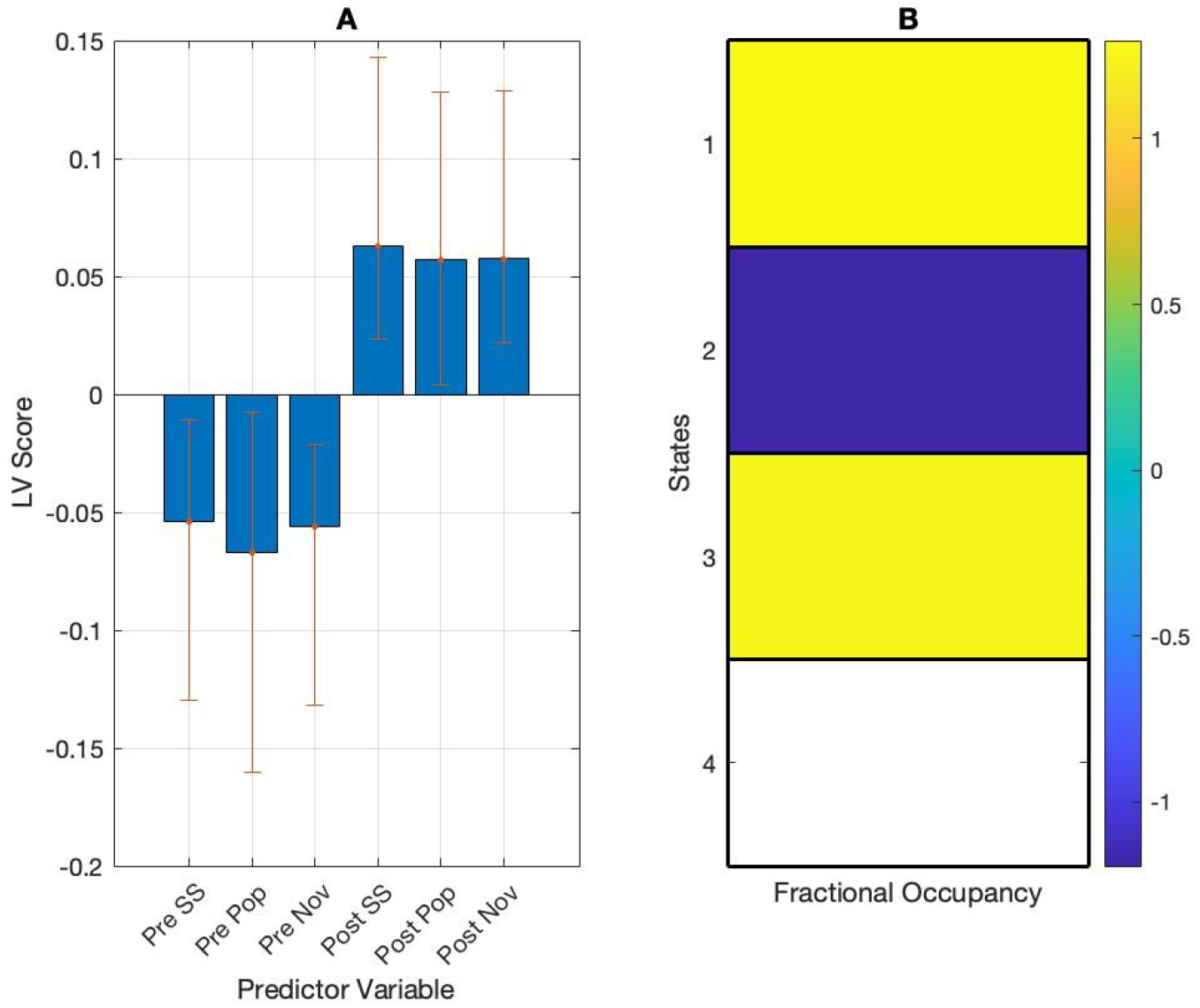
PLS results showing an intervention effect in music listening. (A) PLS contrasts in pre- and post-intervention during music listening. Error bars were calculated using bootstrap resampling and reflect the 95% confidence interval. The contrasts show an intervention effect on FO (B). The colour scale represents the bootstrap ratio for each state. Participants pre-MBI have higher fractional occupancy in state 2 (temporal state) while participants post-MBI have higher fractional occupancy in states 1 (medial frontoparietal) and 3 (temporal-mesolimbic). SS = self-selected, Pop = popular, Nov = novel.

**Table 1:** Regions of interest and anatomical labels from HMM analysis. Anatomical labels are based on the work of Uddin et al. (2019).

| State | Main Regions | Anatomical label |
| --- | --- | --- |
| 1 | Bilateral middle-frontal and left temporal regions. Subcortical regions include the bilateral temporal pole, left nucleus accumbens, and right hippocampal body | Medial frontoparietal |
| 2 | Bilateral temporal and frontal regions | Temporal |
| 3 | Bilateral temporal and mesolimbic regions<br>Subcortical regions include the left globus pallidus, left hippocampal body, right putamen, and right hippocampal tail | Temporal mesolimbic |
| 4 | Bilateral superior frontal and middle parietal regions | Frontoparietal |

PLS analysis on the average fractional occupancy data returned no significant differences between pre- and post-intervention data (*p* = 0.32).

An intervention effect was observed when the data were split into stimulus categories. In these results, the pre-MBI condition showed a higher likelihood of transitioning into state 2 and a higher likelihood of persisting in state 2 in experimenter-selected pieces. The post-MBI condition showed a higher likelihood of transitioning from state 2 to state 3 in all stimulus categories (Figure 3).

**Figure 3:**
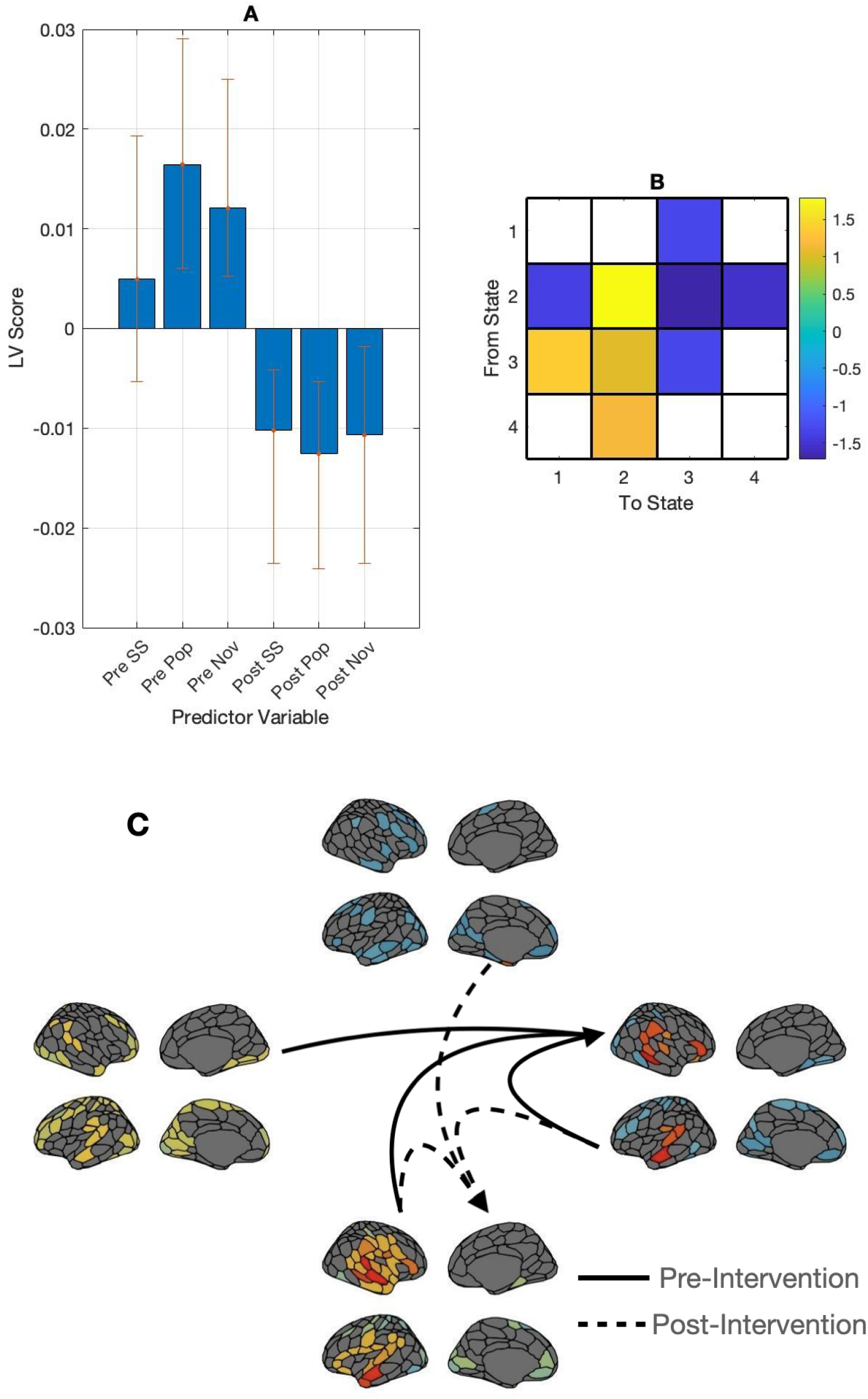
PLS results showing an intervention effect in music listening in transitional probability matrices. (A) PLS contrasts between groups in stimulus categories and transitional probabilities (SS = self-selected, Pop = popular, Nov = novel). Error bars were calculated using bootstrap resampling and reflect the 95% confidence interval. The contrasts (B) show an intervention effect on transitional probability (TP), illustrated in panel C. Panel C shows the between-network TP with solid lines representing self-selected music and dashed lines representing experimenter-selected music. Participants pre-MBI have are more likely to transition into state 2 (temporal) and to stay in this state during experimenter-selected music while participants post-MBI have are more likely to transition from state 2 (temporal) to state 3 (temporal-mesolimbic) for all stimulus categories.

PLS analysis on the averaged pre- and post-MBI transitional probability matrices did not return any significant LVs (*p* = 0.31).

We ran a within-group PLS analysis on the relation between age and HMM features to control for possible age effects in these results. These analyses returned no significant results indicating no age effects on fractional occupancy or transitional probabilities within this sample.

### Effects of liking and familiarity on brain measures

We next correlated participants’ fractional occupancy and vectorized transitional probability ratings with their piece-wise liking and familiarity ratings. PLS analysis returned no significant LVs for fractional occupancy or transitional probability data.

### Behavioural ratings

Behaviourally, liking and familiarity were not significantly different post-MBI, nor were they significantly differently correlated pre- or post-MBI (*p* = 0.89).

## Discussion

In this study, we tested brain state metrics extracted with HMM for intervention effects in healthy older adults who had undergone an eight-week music-based intervention (MBI), finding evidence of an intervention effect following the MBI. Quinci and colleagues found increased functional connectivity between seeds in the auditory network and the medial prefrontal cortex in these data following the MBI (Quinci et al., 2022). The present findings complement those results in the network domain with a few key differences. In the present study, we estimated functional connectivity states from timeseries data using HMM. Working in this state space limits the conclusions we can draw from changes in functional connectivity - when the smallest unit of measurement is a state, the connection weights between regions are fixed. This allows us instead to examine the temporal properties between the states, namely, how often each state is visited and the likelihood of transitioning between each state. While Quinci and colleagues found increased functional connectivity between key seed regions (2022), we here add temporal detail showing that the increased connectivity may be due to more frequent visits and/or a more well-established path to a state containing auditory and reward regions.

Pre-MBI, participants showed higher fractional occupancy and a greater likelihood of transitioning to and persisting in the temporal state in experimenter-selected music. This finding is consistent with results presented in our previous work (Faber et al., 2023), and shows participants occupying the temporal state most often during experimenter-selected music. In our previous study (Faber et al., 2023), older adults showed higher occupancy in the temporal-mesolimbic state compared to younger adults in experimenter-selected music; we now know that this effect is more pronounced post-MBI. Post-MBI, transitioning into the temporal-mesolimbic state was higher than pre-MBI, and transition to, and state persistence in the auditory state was lower following the MBI. These results indicate that the temporal state may become less engaged in favour of the temporal-mesolimbic state following the MBI, similar to the dedifferentiation-like effects seen in the previous study but with an interesting twist: this effect is malleable over time, especially with the intervention.

Numerous studies have focused on dedifferentiation as an age-related process (see Koen and Rugg, 2019), with cross-sectional and longitudinal studies showing effects emerging with age in rest and task experiments (Malagursky et al., 2020; Grady et al., 2016). Dedifferentiation does not robustly predict poor cognition (Koen & Rugg, 2019), and may serve as a beneficial adaptation to age-related perturbations (Rieck et al., 2017; Rieck et al., 2021). With this in mind, the post-intervention increases in fractional occupancy and transitional probabilities involving the temporal-mesolimbic state may be a positive adaptation of the brain in response to music listening. The recruitment of auditory-reward regions where previously a network of auditory-only regions would be employed indicates music, in general, is more rewarding post-MBI.

Though test-retest effects cannot be discounted entirely, the lack of accompanying increases in liking and familiarity ratings suggests that the differences in network activity in the post-MBI group cannot be attributed simply to a greater familiarity with the stimuli. Similarly, the relatively short intervention period renders longitudinal aging-related effects unlikely. Rather, these results suggest a true intervention effect. This is an exciting possibility for clinical research as it suggests that music-based interventions can strengthen the activity of the auditory reward system (or lower its threshold for activation). If music listening stimulates more auditory-reward activity with regular listening in healthy older adults, what results and/or benefits could we expect in clinical populations, namely those with neurodegeneration?

Though these findings are promising, they are not without limitations. The effects we observed, while statistically significant, were calculated on a small sample without a control group. These findings show a dedifferentiation-type pattern where all types of music post-intervention preferentially involve auditory reward regions, but more work is needed, including data from younger adults to establish the wider applicability of these findings. The age range of participants in this sample was broad and while it did not significantly affect these results, a larger sample size may provide enough variability to observe cohort effects within the older adult sample.

These findings add promising information to the growing research on the relationship between brain network dynamics and music-based intervention. Research studies and popular media have shown that music continues to engage individuals with neurodegeneration (Cuddy & Duffin, 2005; Särkämö et al., 2014, Satoh et al., 2015; Baird et al., 2020; Rossato-Bennett, 2014), and from here, several exciting avenues emerge. We have shown changes in state connectivity in a healthy older adult population following a music-based intervention, which will prove useful in further work with clinical populations. This healthy baseline provides a starting point for music-based work with individuals with neurodegeneration. Will these individuals show a similar pattern of auditory reward network activity during music listening? Will these pathways change with music-based intervention, and if so, will it mirror the changes seen in adults without neurodegeneration? Music is unique in its ability to remain engaging and accessible to individuals with neurodegeneration, and has immense potential to teach us about how the brain ages and adapts to neurodegeneration. It is our hope that these findings spark new collaborations between clinicians, researchers, and patient groups striving to better understand the lived experiences of individuals with dementia and the role that music can play as a therapy and as an investigative tool.

